# Evaluation of toxicological mechanisms of ochratoxin-A in human primary proximal tubule epithelial cells

**DOI:** 10.1101/2025.08.29.673189

**Authors:** Anish Mahadeo, Theo K. Bammler, James MacDonald, Angela R. Zheng, Catherine K. Yeung, Jonathan Himmelfarb, Edward J. Kelly

## Abstract

Ochratoxin-A (OTA) is a ubiquitous mycotoxin contaminant in food products and a known nephrotoxin. OTA is hypothesized to be a potential environmental agent causing chronic kidney disease of unknown etiology (CKDu), however the mechanism of OTA toxicity in the human kidney remains elusive. This study aims to elucidate OTA-induced molecular toxicological pathways using primary human proximal tubule epithelial cells (PTECs). We demonstrated that exposure to OTA (10 μM) induces over 7000 differentially expressed genes, including key regulators of mitochondrial fission and fusion. This was confirmed at the cellular level by confocal microscopy, where a breakdown of the mitochondrial network was observed at 100 nM OTA. Crucially, OTA was found to significantly induce reactive oxygen species (mROS) and inhibit basal mitochondrial oxidative phosphorylation as well as glycolysis through measurements of oxygen consumption rate and extracellular acidification, indicating reduced cellular energetics and mitochondrial toxicity. The previously reported downregulation of NRF2 target antioxidant response elements was not able to be recovered with co-administration of NRF2 agonists, sulforaphane or tert-butylhydroquinone, suggesting a possible mechanism of inhibition of NRF2 nuclear translocation or DNA binding. In conclusion, we demonstrate that OTA induces oxidative stress, mitochondrial dysfunction, and reduced ATP production, leading to a senescent-like state in PTECs characteristic of renal disease progression. These findings provide insight into early toxicological endpoints induced by OTA which have been established as pathophysiological changes involved in chronic kidney disease.

**Highlights:**

- OTA exposure suppresses oxidative phosphorylation and glycolysis
- OTA exposure disrupts mitochondrial fission and fusion
- OTA induced mitochondrial ROS
- Early mitochondrial stress may underlie OTA-associated CKDu risk

## 1. Introduction

Ochratoxin-A (OTA) is one of the most abundant mycotoxins found in food products. While the fungal genera (*Aspergillus* and *Penicillium)* that produce this mycotoxin grow well in tropical environments, they can be found ubiquitously around the world, particularly in areas where crop fields and food storage facilities have improper environmental controls. Plant-based products, especially grains, are most susceptible to OTA contamination and include foods such as wheat, rice, corn, coffee, beer and wine (Malir et al. 2016). OTA is classified by the International Agency for Research on Cancer (IARC) as a Group 2B carcinogen, and is a known nephrotoxin, hepatotoxin and neurotoxin (de Groene et al. 1996). As a nephrotoxin, it has been hypothesized that OTA is linked to unexplained instances of chronic kidney disease in endemic regions such as Sri Lanka and Tunisia due to higher concentrations of OTA found in patient blood and urine samples, as well as household food products from endemic regions (Desalegn et al. 2011; Maaroufi et al. 1995). However, conflicting findings have been reported in other regions of the world, where clinical OTA concentrations do not seem to be correlated with renal disease (Domijan et al. 1999; Fazekas et al. 2005; Fuchs and Peraica 2005; Kosicki et al. 2021; Palli et al. 1999).

This unexplained chronic kidney disease, known as chronic kidney disease of unknown etiology (CKDu), disproportionately affects tropical, low-to-middle income countries such as India, Sri Lanka and the Central American region. In many of these regions, CKDu accounts for 50-70% of all reported chronic kidney disease cases (Jayasekara et al. 2019). The pathophysiology of CKDu differs from chronic kidney disease in that patients display a high degree of proximal tubulointerstitial nephritis rather than glomerulosclerosis. In the past two decades, research into this condition has suggested that CKDu may be the result of a multifaceted set of environmental agents, such as heavy metals, pesticides, herbicides, and mycotoxins (Paidi et al. 2021; Rajapaksha et al. 2021). However, these studies have been correlative and as of yet, no strong mechanistic studies have linked any of these agents to the progression of CKDu. This present study investigates one agent, OTA, at a molecular and cellular level in order to determine its renal toxicological potential.

In previous rat models, OTA was shown to accumulate in the kidney at levels 12-fold higher than that of plasma after intraperitoneal injection (Schwerdt et al. 1996). Additionally, *in vitro* transport studies indicate that OTA is transported with high-affinity from the bloodstream into the proximal tubule by organic-anion transporters 1 and 3 (OAT1/3) but is transported with low affinity into the urinary filtrate by efflux transporters. It has also been shown that OTA is reabsorbed from the urinary filtrate with high-affinity by organic anion transporter 4 (OAT4), which, taken together indicates a high potential for accumulation in the proximal tubule (Anzai et al. 2010; Jutabha et al. 2011; Studer-Rohr et al. 2000). Several hypotheses of toxicological mechanisms have been proposed for OTA, yet predominant pathways remain unclear. Several studies indicate the potential of OTA to produce oxidative stress *in vitro* and in some *in vivo* rat models by superoxide and hydroxyl radical generation (Babayan et al. 2020; Gautier et al. 2001; Ramyaa and Padma 2013; Schaaf et al. 2002). Research indicates the downregulation of nuclear erythroid-related factor 2 (NRF2) target antioxidant response genes in response to OTA treatment as a major cause of oxidative stress (He and Ma 2009; Loboda et al. 2017; O’Mealey et al. 2017; Shin et al. 2019). Genotoxicity and adduct formation have also been reported as toxicological mechanisms via formation of both a phenoxyl radical and CYP450 bioactivation into a quinone metabolite (Babayan et al. 2020; de Groene et al. 1996; Mantle et al. 2010). Other studies indicate the potential of OTA to inhibit protein synthesis and cause epigenetic changes leading to disease states in different tissues (Babayan et al. 2020; Creppy et al. 1984). Furthermore, a few studies have indicated that OTA induces mitochondrial reactive oxygen species (mROS) production and disruption of oxidative phosphorylation, promoting tumorigenesis and neuronal disease (Chung et al. 2019; Park et al. 2019).

Previous research on OTA toxicology has suggested numerous biochemical pathways leading to cytotoxicity, but few studies have focused on mechanisms in the proximal tubule. Here, we utilized human primary proximal tubule epithelial cells (PTECs) to build upon previous findings from our laboratory and others to address this issue. Clinical data has indicated a varying range of OTA in human samples between healthy individuals and patients with an estimated plasma concentration of 0.7 to 7.8 ng/mL (1.7 to 19 nM), with CKD patients in Tunisia reporting up to 815 nM plasma OTA concentration (Abid et al. 2003; Coronel et al. 2011; Maaroufi et al. 1995; Palli et al. 1999). As this renal site may be the target of OTA accumulation, the present study aimed to elucidate biochemical mechanisms of toxicity induced by OTA in the proximal tubule at physiologically relevant concentrations from 1 nM to 10 μM.

## 2. Materials and Methods

### 2.1 Cell culture

Human kidneys used to isolate cells in this study were obtained from Novabiosis (Durham, NC), an organ procurement organization of discarded deceased donor tissue. Demographics collected from de-identified electronic medical records are listed in Supplemental Table 1. Isolation of primary human proximal tubule epithelial cells was performed as described previously (Van Ness et al. 2017) and stored in liquid nitrogen until experiments were ready to perform. PTECs were thawed and cultured in DMEM (Gibco, Waltham, MA, 1130-032) with 1x insulin-transferrin-selenium-sodium pyruvate (ITS-A, Gibco, 51300044), 50 nM hydrocortisone (Sigma, H6909) and 1x Antimicrobic-Antimycotic (Gibco, 2520056) until confluence. Cells were passaged with 2-minute 0.05% trypsin-EDTA (Gibco, 25200056) exposure and neutralized with defined trypsin inhibitor (Gibco, R007100), then centrifuged at 400g for 5 minutes, and finally re-plated for experiments with the above supplemented DMEM solution. Cells were incubated at 37°C and 5% CO2. Isolation of mRNA and mRNA-Sequencing

PTECs of donor ID PT6 and PT10 (Passage 2), in triplicate, were plated in two 24-well plates (Corning, Corning, NY, 3548) in supplemented DMEM medium and grown to confluence.

Treatment group samples were pre-treated for 48 hours with DMEM medium and either sulforaphane (Sigma, St. Louis, MO, S4441), or tBHQ (Sigma, 112941) and controls were treated with supplemented DMEM medium. Then, plates were treated for 72 hours with ochratoxin-A (Sigma, O1877) in either supplemented DMEM, supplemented DMEM + sulforaphane, or supplemented DMEM + tBHQ. Following 72-hour incubation, cells were lysed with Buffer RLT (Qiagen, Hilden, DE, 74004) and mRNA extraction and sequencing was performed by Novogene (Sacramento, CA). Read quality was assessed using FASTQC (https://www.bioinformatics.babraham.ac.uk/projects/fastqc/), and then reads were aligned to the Ensembl GRCh38 patch release 27 genome using the HISAT2 aligner(Kim et al. 2019). Reads were imported into R, summarized as reads/gene using the Bioconductor Rsubread package(Liao et al. 2019), and then analyzed using the limma-voom pipeline(Law et al. 2014). An additional fold change criterion was included in the statistical analysis to restrict to those genes with likely biologically meaningful changes in expression (McCarthy and Smyth 2009). Statistical significance was determined using the Benjamini-Hochberg procedure.

### 2.2 Mitochondrial and glycolytic stress analysis

A Seahorse XF96 Analyzer (Agilent, Santa Clara, CA) was used to determine oxygen consumption rate (OCR) and extracellular acidification rate (ECAR). The optimal concentration of cells that produced reliable responses to assay reagents in the XF-96 well plate was experimentally determined to be 24,000 cells per well. In this experiment, PT11, PT23, and PT21 (Passage 2, 3 and 2 respectively) were used. Cells were allowed to adhere to the plate before being treated with OTA for 24 hours. The Seahorse XF Mito Tox Assay kit (Agilent, 103595-100) and glycolytic stress test kit (Agilent, 103020-100) were performed 24 at this point as per manufacturer’s protocol. 2 μM Oligomycin, 3 μM FCCP, and 1 μM Rotenone/Antimycin A were used in injections. Prior to the assay, cells were incubated in plain XF DMEM media (Agilent, 103575-100) without CO_2_ for 1 hour and cell health was assessed by light microscopy. For the glycolytic stress test, 50 mM glucose was injected at 20 minutes into the assay, 3 μM FCCP was injected at 40 minutes, and 100 mM 2-deoxyglucose (2-DG) was injected at 60 minutes. Data was analyzed using Seahorse Wave Desktop software 2.6.1 (Agilent). Significance was determined in GraphPad Prism 10 using an ordinary one-way ANOVA with Dunnet’s post hoc test.

### 2.3 MitoSOX Assay and MitoTracker Red Staining

For the MitoSOX experiment, three donors of PTECs (PT11 passage 2, PT23 passage 3, PT21 passage 3) were grown to confluency in 24-well plates. Cells were treated with control media, 10 μM ochratoxin-A, or 25 μM menadione (Sigma, M5625) for 24 hours with five technical replicates per donor, per treatment. Cells were then washed three times with warm PBS prior to addition of 5 μM of MitoSOX green reagent (optimized concentration, ThermoFisher, Waltham, MA, M36005), incubated at 37°C and 5% CO_2_ for 30 minutes. Cells were then washed again three times with warm PBS and immediately imaged using a Nikon Ti-Eclipse fluorescent microscope. Treatments in each well were coded and administered by an independent experimenter so imaging and analysis could be done in a blinded manner. Image analysis was performed in Python using the Scikit-image package for background subtraction, binarizing and intensity quantification. Statistics were performed using GraphPad Prism 10 with a one-way ordinary ANOVA test.

PT11 (passage 2) cells were grown to confluency in two 24-well plates. Cells were then treated with control media or 10 μM OTA. One plate was transferred to a hypoxia incubator with oxygen levels maintained at 2% with 5% carbon dioxide. The remaining plate was incubated at normal conditions described above. After 24 hours of incubation, plates were removed from the incubators and media was replaced with pre-warmed (37°C) media containing 500 nM MitoTracker Red FM (ThermoFisher, M22425). Plates were returned to their respective incubators for 30 minutes. Cells were then washed with warm PBS and fixed with 4% formaldehyde for 15 minutes. Plates were then immediately imaged using a Nikon Ti-Eclipse fluorescent microscope.

### 2.4 Mitochondrial labeling and confocal imaging

Human primary proximal tubule cells (PT6, PT10, PT23, passage 2) were grown to 50% confluence in a T25 plate. At this point, cell media as described before was made with 10 µg/mL polybrene. Cells were transduced with pCT-COX8-GFP (developed by Drs. Lans Taylor and Larry Vernetti, University of Pittsburgh) at a multiplicity of infection of 10:1 to cell concentration. Cells were grown to confluency and checked for reporter expression by fluorescent microscopy. These cells were plated into Ibidi u-Slide 8 well microscopy slides (Ibidi, Gräfelfing, DE, 80826), then treated with indicated OTA concentrations with or without 1 μM cobalt chloride, as well as 100 μM Apelin-13 (Sigma, A6469). Imaging was done using a Nikon A1R confocal microscope with 60x oil immersion lens, 2x line averaging, 1024 by 1024 resolution, and 3% laser power. Image analysis was done in ImageJ using the MiNA 2.0.0 ImageJ plugin. Mitochondrial footprint was calculated by dividing mitochondrial area (MiNA output) by cell area (measured in ImageJ). One-way ANOVA with Sidak’s multiple comparison as well as graph generation from the MiNA output was done in GraphPad Prism 10 (Dotmatics, Boston, MA). Treatments in each well were coded and administered by an independent experimenter so imaging and analysis could be done in a blinded manner.

### 2.5 Cytokine Quantification

Cytokine quantification reported in this study was performed by ELISA. PTECs were plated in 24-well plates at 250,000 cells/mL and allowed to reach confluency, examined by light microscopy. In this experiment, PT11, PT21, PT23 were used. Cells were then treated with 10 μM OTA, 20 μM 2’3’-cyclic GMP-AMP (cGAMP, Invivogen, San Diego, CA, tlrl-nacga23), with or without H-151 (Invivogen, inh-h151) for 24 hours. Cell media was then collected to be analyzed by ELISA. IL-6 and CXCL6 ELISA kits were acquired from R&D systems (Catalog # DY206-05 and DY333, respectively) and experiments were conducted following manufacturer’s protocol. Dilution of samples was experimentally determined to be 1:10 for IL-6 and 1:2 for CXCL6. Fluorescence was measured using a Tecan Spark multimode microplate reader (Tecan Trading AG, Männedorf, Switzerland). Data was normalized to protein level by BCA assay (Thermo, 23225) and analyzed by GraphPad Prism 10 (Dotmatics) using the Kruskal-Wallis with Dunn’s post-hoc multiple comparison test.

### 2.6 Establishment of Microphysiological Systems (MPS) and Immunocytochemistry

Nortis ParVivo triple channel MPS devices were prepared as described previously(Chalker et al. 2025; Van Ness et al. 2017). Chambers were filled with 6 mg/mL rat tail collagen 1 (Ibidi, 50204) and allowed to polymerize overnight at room temperature. Cell media was then perfused through the devices at 1 µL/min. Prior to cell seeding, 5 µg/mL mouse collagen IV (Corning, 47743) solution was injected into the tubules and incubated for 30 minutes. Tubules were then washed with media perfusion for 1 hour at 1 µL/min. PTECs (PT10, passage 2) were then seeded into the devices at 10 million cells per milliliter and allowed to attach for 3 hours. Media perfusion was then initiated and tubules were allowed to grow until confluency, verified by light microscopy. Upon tubule formation, tubules were perfused with media or media containing 10 μM OTA, 1 μM cobalt chloride or both for 48 hours (n=1). For immunocytochemistry, tubules were first fixed by perfusion of 4% formaldehyde solution for 30 minutes, followed by a 30-minute PBS wash. Immunocytochemistry perfusions were performed at 10 µL/min. Tubules were then perfused for 2 hours with PTB solution: PBS solution containing 50mg/mL bovine serum albumin and 0.1% Triton X-100 (Sigma-Aldritch, 9036-19-5). PTB solution containing a 1:100 dilution of anti-HIF1 alpha antibody (Abcam, ab51608) was then perfused for 1 hour and incubated without flow for another hour at room temperature. Following a 2 hour wash with PBS, PTB solution containing a 1:1000 dilution of goat anti-rabbit Alexa Fluor 488 was perfused for 1 hour. Secondary antibody was then washed out with PBS perfusion for 2 hours. PBS solution containing 1 µg/mL Hoechst nuclear stain (ThermoFisher, 62249) was then perfused for 20 minutes. Imaging was performed using a Nikon Ti-Eclipse fluorescent microscope.

## 3. Results

### 3.1 OTA dysregulates mitochondrial homeostasis at both the transcriptional and cell physiological level

Whole mRNA sequencing analysis of human primary PTECs treated with OTA revealed significant differentially expressed genes involved mitochondrial fission, fusion and mitophagy/apoptosis following 72-hour OTA administration (Figure 1A) in two cell donors. To investigate this phenomenon at the cellular level, we transduced PTECs with a Lenti virus containing cytochrome-C-oxidase subunit 8 – GFP (COX8-GFP) to label mitochondrial inner membranes and exposed these cells to OTA concentrations (100nM or10 μM) with or without 1 μM cobalt chloride to simulate low oxygen conditions for 24 hours. Confocal microscopy revealed severe aberrant mitochondrial phenotypes even at 100 nM concentrations of OTA. Figure 1B shows fragmentation of the mitochondrial network at 100 nM OTA exposure, and severe breakdown of the network and hyperfused mitochondrial structures with 10 μM OTA and the positive control, 1 μM cobalt chloride. Cobalt chloride was selected as a positive control as it is a DRP1-dependent mitochondrial fragmentation inducer as well as a hypoxia-mimetic agent, which simulates oxygen conditions in the renal cortex (Zheng et al. 2020). Reduction in mitochondrial signal was also observed with 10 μM OTA exposure in PTECs cultured under 2% oxygen conditions (Figure S2). Apelin-13, a regulator of mitochondrial homeostasis via Sirtuin 3 (SIRT3) was also administered to attenuate network disruption. Mitochondrial geometric analysis in ImageJ was used to characterize mitochondrial network structures in these treatment conditions (Figure 1D), and this revealed a significant breakdown of mitochondrial area (left) in one donor, network branching complexity (middle), and the ratio of fragmented structures to more complex structures (right) with all concentrations of OTA. Apelin-13 co-administration successfully attenuated the morphological changes in mitochondrial in all three of these parameters, except for mitochondrial footprint in PT10 and branches per network in PT6. Additionally, numerous cells treated with OTA at both 100 nM and 10 μM (50% and 53%, respectively) were observed to have hyperfused mitochondria (Figure 1C, Table 1) compared with 10% of cells treated with control and 13% of cells treated with cobalt chloride. This phenomenon was observed to begin after 4 hours of 10 μM OTA exposure, evident after 12 hours in one donor (Supplementary Figure 1, OTA_ROI.avi and control_timelapse.avi) but not in control cells.

**Figure 1.**
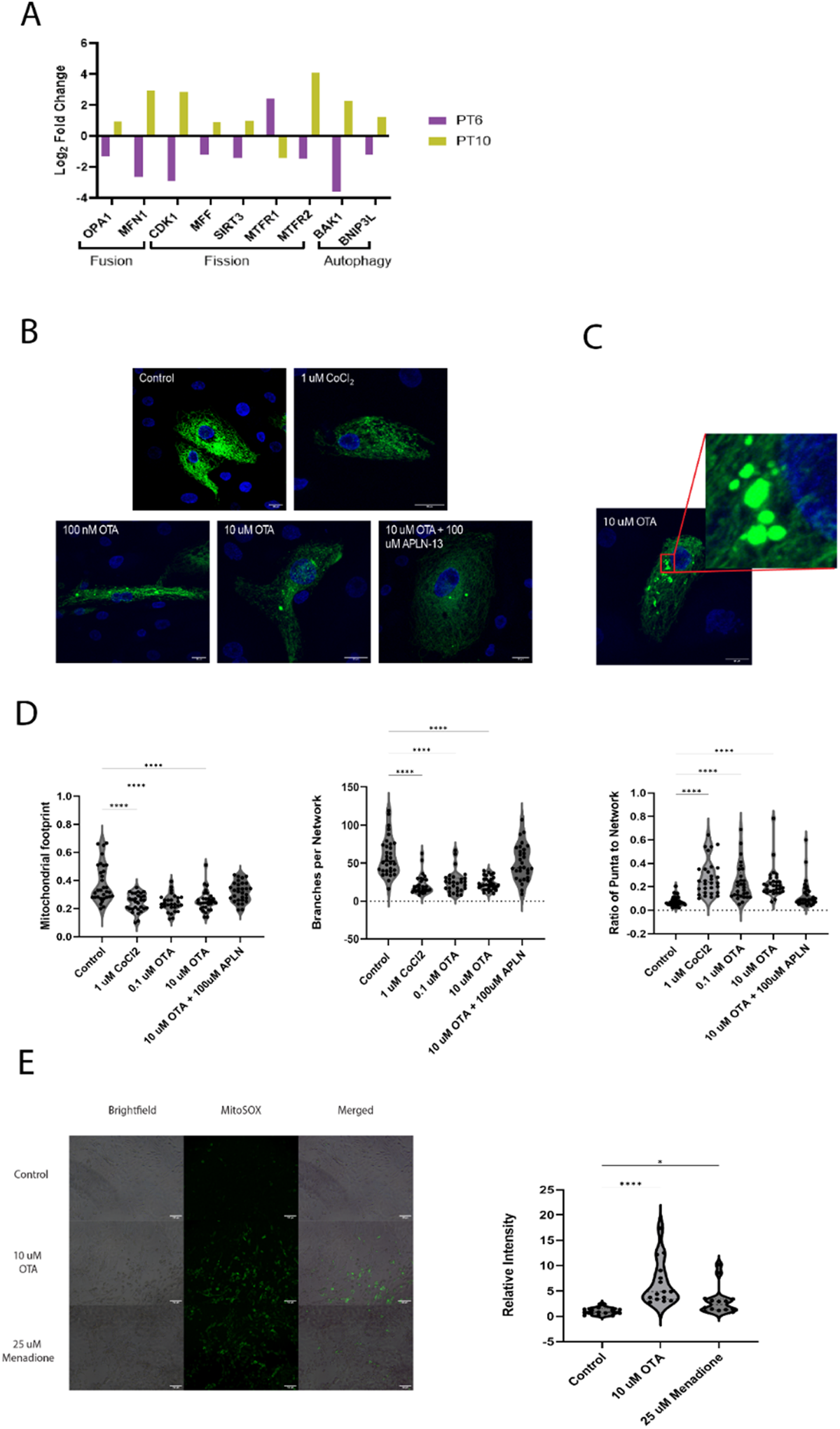
OTA disrupts homeostasis between mitochondrial fission and fusion and induces oxidative stress. A) mRNA expression of differentially expressed genes involved in mitochondrial fission, fusion and mitophagy. Abbreviations: Mitochondrial dynamin like GTPase (OPA1), mitofusin 1 (MFNl), cyclin dependent kinase 1 (CDKl), mitochondrial fission factor (MFF), mitochondrial fission regulator 2 (MTFR2), BLC2 antagonist/killer 1 (BAKl), BLC2 interacting protein 3 like (BNIP3L). **B)** Representative confocal images (60x) of PTECs (PT6, PTl0, PT23) transfected with mitochondrial membrane COX8-GFP (green) as well as Hoechst nuclear stain, treated under different conditions. Scale bar: 20 µm. **C)** Representative image of hyperfused mitochondria {expanded panel) under 10 µMOTA treatment. Scale bar: 20 µm. **D)** Mitochondrial network characterization performed with the MiNA plugin in lmageJ (Valente et al. 2017). Data shows 10 cells from biological donors PT6, PT10 and PT23. **E)** Left panel: Representative images of mitochondrial-specific reactive oxygen species analyzed by fluorescence microscopy with the MitoSOX Green dye under different treatments. Scale bar: 100 µm. Right panel: Relative fluorescence intensity analysis. Three biological replicates were used (n=3; PT11, PT21, PT23) with 5 culture wells each. * indicates p<0.05, ** indicates p<0.005, *** indicates p<0.0005, **** indicates

**Table 1.** OTA exposure induces the formation of hyperfused mitochondria in PTECs.

| **Treatment** | **% of Cells with Hyperfused Mitochondria\*** |
| --- | --- |
| Control Media | 10 |
| 100 nM OTA | 50 |
| 10 μM OTA | 53 |
| 10 μM OTA + 100 μM APLN-13 | 33 |
| 1 μM CoCl<sub>2</sub> | 13 |

*\*Percentages of cells across three cell donors (PT10, PT6 and PT23) observed to contain hyperfused mitochondria observed with confocal microscopy. An example of hyperfused mitochondria is reported in Figure 2C.*

### 3.2 Ochratoxin-A disrupts mitochondrial function and cellular metabolism

Given that OTA was shown to alter the structure of mitochondrial networks in PTECs, we next analyzed the effects of OTA on mitochondrial function, specifically oxidative phosphorylation and ATP generation by measurements taken with a Seahorse XF Flux Analyzer. Indeed, OTA was shown to significantly reduce basal oxidative phosphorylation (p < 0.0005), maximal respiration (p < 0.05) and ATP production (p < 0.0005) by 30% at concentrations as low as 10 nM and by 41% with 10 μM OTA (Figure 2A). We next hypothesized that with oxidative phosphorylation reduction, there may be an increase in cellular glycolysis to compensate for the loss of ATP. We analyzed glycolysis by measuring extracellular acidification rate (ECAR). Surprisingly, no change in glycolytic rate was observed up to 1 μM OTA, where basal glycolysis was decreased by 33% (Figure 2B). There was a modest but significant decrease in glycolytic capacity at 10 μM OTA. The reduction in both oxidative phosphorylation and glycolysis is plotted (Figure 2C), depicting a shift in metabolic phenotype from “energetic” to “quiescent.” These experiments indicate that OTA reduces not just oxidative phosphorylation, but overall cellular metabolism.

**Figure 2).**
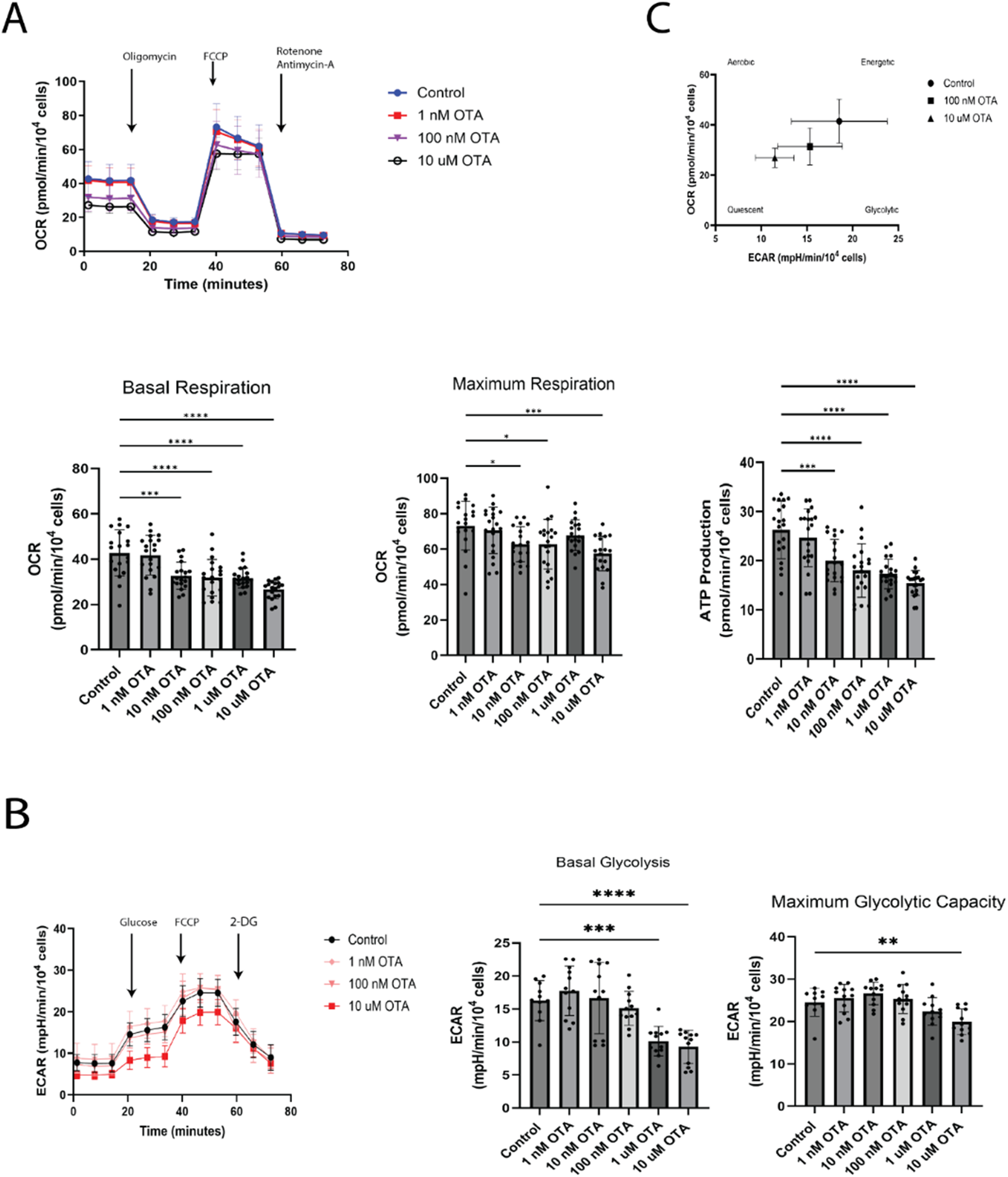
OTA reduces mitochondrial oxidative phosphorylation and glycolysis. **A)** Measurement of Oxygen Consumption Rate (OCR) of PTECs (PT21, PT23, PT6) treated with increasing dose (1 nM to 10 µM) of OTA for 24 hours compared to control cells. Top panel: time-course of OCR with administration of oligomycin, carbonyl cyanide-p-trifluoromethoxyphenylhydrazone (FCCP), and rotenone with antimycin­A. Bottom panels: Basal respiration, maximal respiration and ATP production. **B)** Measurement of glycolysis by extracellular acidification rate (ECAR) with increasing dose of OTA compared with control. left: time-course of ECAR with administration of glucose, FCCP and 2-deoxyglucose (2-DG). Basal glycolysis and maximum glycolysis measurements are shown in the middle and right, respectively. **C)** Cellular energy phenotype profile of cells treated with increasing concentrations of OTA compared with control. Experiments were conducted in biological triplicate. * indicates p<0.05, ** indicates p<0.005, ***

### 3.3 Downregulation of NRF2-target genes due to OTA were not ameliorated by NRF2 agonists

We have previously shown that OTA administration to PTECs leads to the downregulation of numerous NRF2 target mRNA transcripts, but the mechanism is unknown (Imaoka et al. 2020).

While the exact mechanism of OTA-dependent inhibition of target genes is not known, it has been hypothesized that OTA blocks the liberation of NRF2 from KEAP1 which prevents the nuclear translocation of NRF2, or it may inhibit DNA binding (Limonciel and Jennings 2014). First, we confirmed that NRF2 target genes would be dysregulated in PTECs after 72-hour exposure to 10 μM OTA. Whole mRNA sequencing revealed numerous dysregulated NRF2 target genes (WikiPathway 2884) in two PTEC donors. Interestingly, we found that different sets of these NRF2 targets were upregulated and downregulated in the two donors (Figure 3A), but nonetheless numerous NRF2 targets were significantly downregulated in each donor. We then investigated if co-administration of NRF2 activators, sulforaphane (SFN) and tert-butylhydroquinone (tBHQ) could revert the OTA-induced NRF2-target mRNA downregulation. These activators release NRF2 from its cytosolic binding/ubiquitin ligase complex KEAP1/CUL3. PTECs were pre-treated with either SFN (20 μM) or tBHQ (5 μM) for 48 hours before administration of OTA (10 μM) for 72 hours. The number of significant differentially expressed genes for each treatment comparison are shown in Table 2. 10 μM OTA with or without activators induced differential expression of more than 7000 genes (FDR<0.05) in both donors. tBHQ exposure significantly induced numerous NRF2-target genes in one donor including HMOX1 and NQO1 (FDR < 0.0005, Figure 3B), and while not significant, an elevated trend in expression for numerous NRF2 targets were observed with exposure to either activator in both cell donors. In every case OTA + tBHQ exposure significantly reverted the induction observed with tBHQ treatment alone (Supplementary Materials, PT6_LogCPM.csv and PT10_LogCPM.csv). Figure 3C compares the Log_2_ fold change of the treatments versus control for numerous NRF2 targets, indicating the lack of efficacy of the NRF2 activators to ameliorate the dysregulation induced by OTA. While some differences are seen between OTA treatment and OTA plus the activators (e.g. PT6, AKR1C2), statistical comparison of OTA alone versus OTA with activators indicated no significance in any of the NRF2 targets. This demonstrates that OTA does not hinder the release of NRF2 from KEAP1 (Figure 3D).

**Figure 3.**
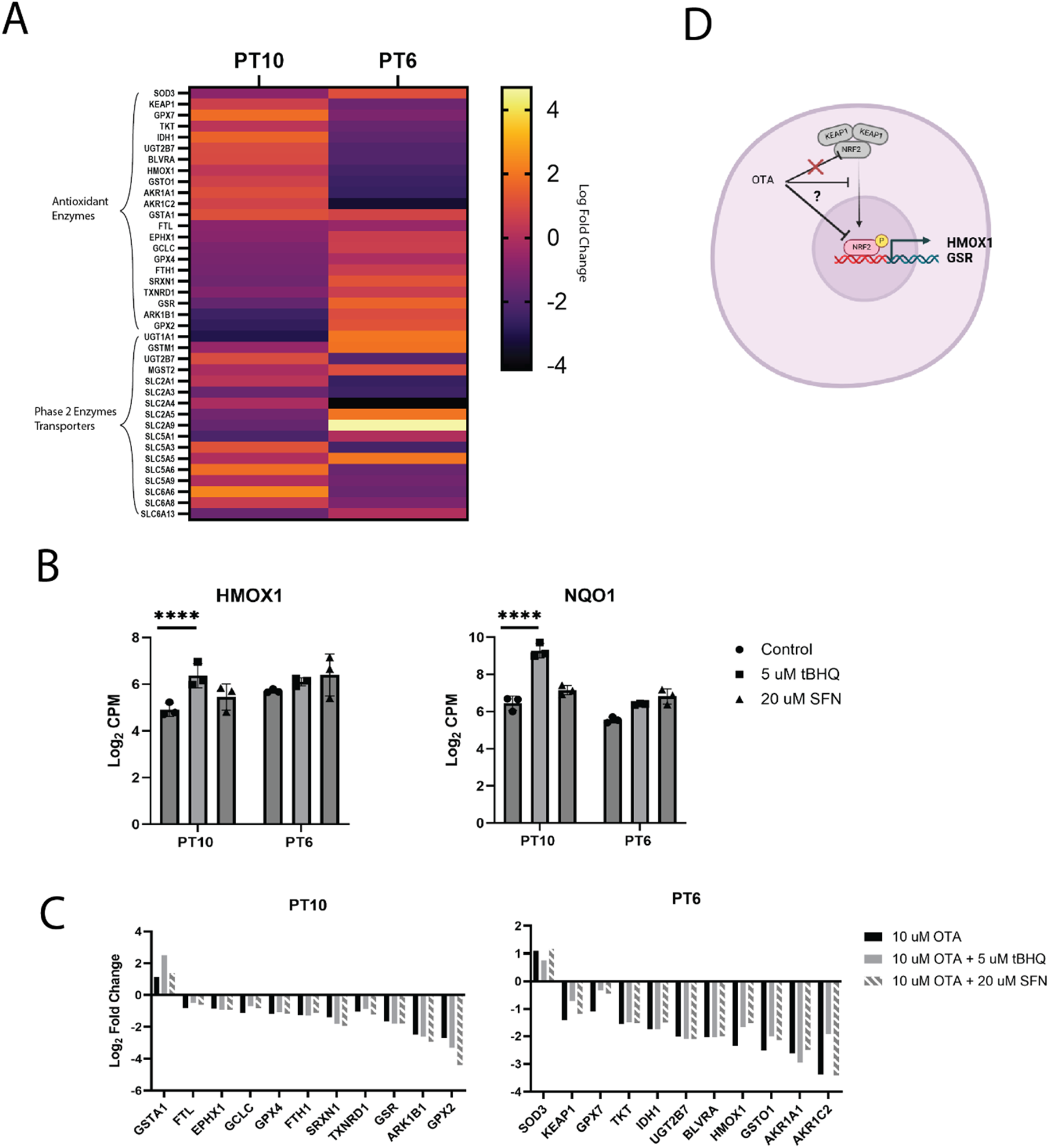
NRF2-target downregulation is not due to direct interaction of OTA with the KEAPl-NRF2 complex. **A)** RNA-seq heatmap showing significant dysregulation of NRF2-target AREs (WikiPathways, WP2884) due to 10 µMOTA treatment for 72 hours. Data is shown for two donors of primary human proximal tubule epithelial cells, n=3 in each case with the average Log_2_ fold change depicted by color. **B)** NRF2-target genes, HMOX1 and NQO1 were significantly upregulated in response to 5 µM tert­butylhydroquinone (tBHQ) exposure(**** indicates FDR< 0.00005) for 72 hours. While not significant, Log_2_ counts-per-million (CPM) expression values were elevated with 72-hour exposure to 20 µM sulforaphane (SFN). **C)** Log_2_ fold change versus control for dysregulated genes after 72-hour exposure to 10 µMOTA, 10 µM OTA+ 5 µM tBHQ, 10 µM OTA+ 20 µM SFN for the two cell donors. All genes shown in panels A and B were significantly dysregulated (adjusted FDR<0.05) versus control and determined by Benjamini-Hochberg procedure. **D)** Graphical depiction of findings in relation to the

**Table 2.** Number of significant differentially expressed genes from control with different treatment conditions.

| **Treatment** | **PT10** | **PT6** |
| --- | --- | --- |
| 10 μM OTA vs. Control | 7192 | 7799 |
| 20 μM SFN vs. Control | 156 | 17 |
| 5 μM tBHQ vs. Control | 1017 | 59 |
| 20 μM SFN + 10 μM OTA vs. Control | 7106 | 7522 |
| 5 μM tBHQ + 10 μM OTA vs. Control | 7276 | 7633 |
| 20 μM SFN + 10 μM OTA vs. 10 μM OTA | 0 | 0 |
| 5 μM tBHQ + 10 μM OTA vs. 10 μM OTA | 6 | 0 |

*Significance was determined with a fold change greater than 1.5 at an FDR less than 5%.*

### 3.4 Investigation of pro-inflammatory pathways induced by cytosolic mtDNA via cGAS-STING

With evidence supporting mitochondrial disruption due to OTA exposure, we next investigated whether the pro-inflammatory cGAS-STING pathway could be activated due to cytosolic mitochondrial DNA (mtDNA) release. mRNA sequencing suggested the possible involvement of this pathway as numerous C-X-C chemokine ligand family proteins, downstream targets of STING activation, were found to be significantly upregulated in PTECs exposed to 10 μM OTA in one cell donor, PT6 (Figure 4A) and interleukin-6 (IL-6) in both donors. We next examined protein levels of IL-6 and CXCL6 by ELISA in PTECs, using 2’3’ cyclic guanosine monophosphate-adenosine monophosphate (cGAMP) as a positive control for STING activation and H-151 as an inhibitor of cGAS-STING. 24-hour exposure of PTECs to OTA showed a 3-fold increase in IL-6 levels compared with control, however this induction was not successfully reverted with co-treatment with H-151 (Figure 4B). Surprisingly, despite induction of mRNA, CXCL6 levels were slightly reduced with OTA treatment, although not significant, and was still significantly reduced compared to control with H-151 treatment.

**Figure 4.**
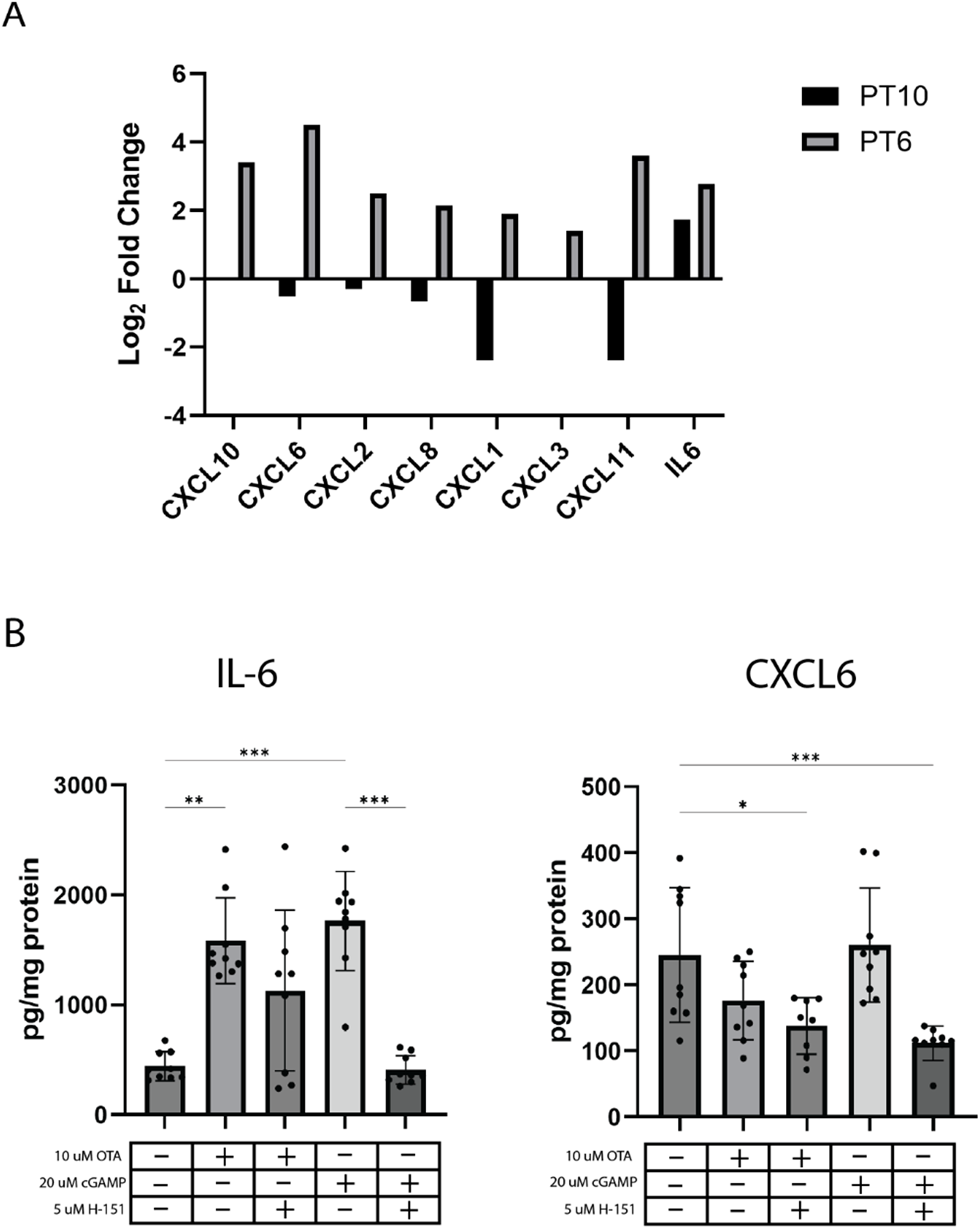
mRNA dysregulation of pro-inflammatory factors may involve complex mechanisms. **A)** mRNA log2 fold-changes of various pro-inflammatory cytokines and chemokines in two biological donors after 72 hour exposure to 10 µMOTA versus control media. **B)** levels of ll-6 andCXCL-6 in PTECs under treatments of OTA or cyclic guanosine monophosphate-adenosine monophosphate (cGAMP) with or without H-151. ELISA experiments were conducted in biological triplicate using cell donors PT6, PT21 and PT23. * indicates p<0.05, ** indicates p<0.005, *** indicates p<0.0005, **** indicates p<0.00005.

## 4. Discussion

The primary goal of this study was to identify important toxicological mechanisms of OTA specifically in the human proximal tubule using primary human proximal tubule epithelial cells.

Mitochondria are very susceptible to oxidative stress; therefore, we hypothesized potential effects of OTA on mitochondrial structure and function. Mitochondrial fusion and fission are finely-tuned processes that are tightly regulated to balance ATP production efficiency with reactive oxygen species production (Adebayo et al. 2021). Treatment of PTECs with 10 μM OTA significantly dysregulated key genes involved in mitochondrial fusion, fission as well as pro-apoptotic proteins (Figure 1A). Significant dysregulation of OPA1 and MFN1, key regulators of mitochondrial fusion (Cipolat et al. 2004) was shown alongside differential regulation of key mitochondrial fission regulators. Differentially expressed genes shown in Figure 2A show activation of both fission and fusion regulators, thus phenotypic analysis and conclusions based off mRNA become difficult. Under normal conditions mitochondria form connective, web-like structures regulated by a balance of biogenesis, fusion and fission events, where shared mtDNA and machinery facilitate efficient oxidative phosphorylation and reduced susceptibility to mitochondrial ROS (Gilkerson et al. 2000; Karbowski and Youle 2003; Skulachev 2001). Breakdown of this network occurs during stress conditions or exposure to mitochondrial toxins, and cisplatin nephrotoxicity has been shown to involve mitochondrial fragmentation and mitophagy (Legros et al. 2002; Zhao et al. 2017). We observed a significant breakdown of this network with 100 nM OTA (Figure 1B, D). Compared to control cells, 100 nM OTA exposure resulted in a 36% decrease in mitochondrial footprint, a 52% reduction in network branching, and a 3-fold increase in the ratio of puncta to network structures, comparable to cobalt chloride (Figure 1D). OTA-induced mitochondrial network degradation (Figure 1D) was successfully attenuated with Apelin-13 co-administration, which has been reported in mice to be an inhibitor of the Mst1-JNK-Drp1 pathway that mediates mitochondrial membrane cleavage (Guo et al. 2024), indicating a potential mechanism of OTA - DRP1 regulation. Interestingly, we also observed an increase in percentage of cells displaying hyperfused mitochondria in response to 100 nM and 10 μM OTA exposure compared with control, an effect that was not observed with cobalt chloride treatment (Table 1). Hyperfused mitochondria have been demonstrated to produce excessive ROS and decreased capacity of oxidative phosphorylation leading to cellular senescence *in vitro* and in rat models of diabetic kidney disease (Yoon et al. 2006). Mitochondria-specific superoxide levels were then assessed by MitoSOX green reagent using 25 μM menadione as a positive control (Loor et al. 2010). 24-hour OTA exposure increased levels of superoxide 6-fold compared to control, compared with menadione which increased levels 3-fold to control (Figure 1E). These results indicate that OTA is compromising mitochondrial dynamics, which is indicative of mitochondrial toxicity and has been implicated as a hallmark of acute kidney injury, chronic kidney disease and cisplatin-induced nephrotoxicity (Fontecha-Barriuso et al. 2022; Tagaya et al. 2022; Zhang et al. 2021).

We next investigated the effect of OTA on oxidative phosphorylation, the primary function of mitochondria. Figure 2A indicates a 34% reduction in basal oxidative phosphorylation was shown with OTA concentrations as low as 10 nM, with 10 μM OTA exposure showing a 42% reduction in OCR. There was also a significant decrease in the maximum rate of oxidative phosphorylation (23%) with 10 nM OTA, decreasing by 27% with 10 μM. ATP production rate was significantly reduced by 30% with 10 nM OTA and up to 55% with 10 μM. Decrease of maximum oxidative phosphorylation is indicative of mitochondrial toxicity as the machinery for ATP production and the ability for cells to respond to a stressor (3 μM FCCP) is reduced. Decreased basal oxygen consumption is indicative of renal hypoxia, and in a microphysiological model of the proximal tubule that we have previously described (Imaoka et al. 2020; Van Ness et al. 2017; Weber et al. 2016), we observed slight stabilization of hypoxia-inducible factor-1 (HIF1α) after 48 hours of exposure to 10 uM OTA, and this effect was potentiated with co-administration of cobalt chloride, shown by a stronger signal than in the individual treatments (Figure S4). As the partial pressure of oxygen in the renal cortex is 2 to 4-fold lower than atmospheric oxygen (Edwards and Kurtcuoglu 2022), this observation in our model supports the potential for OTA to induce hypoxic stress. Interestingly, we also saw a decrease in basal glycolytic rates (extracellular acidification rate, ECAR) only at 1 μM and 10 μM (34% and 38%, respectively) with lower concentrations showing no significant change. Maximum rates of glycolysis only decreased modestly by 20% with 10 μM OTA. Glycolytic enzymes such as glyceraldehyde 3-phosphate dehydrogenase (GAPDH) can be inhibited by ROS, which may explain these results (van der Reest et al. 2018). Figure 2C shows a metabolic phenotype map of control, 100 nM and 10 μM OTA plotted by ECAR and OCR. Compared to control, OTA induces a “quiescent” metabolic phenotype indicated by reduction in both OCR and ECAR. Cellular metabolic quiescence can be induced by stress conditions, where they enter a dormant-like state and conserve energy to combat oxidative stress (Du et al. 2023). This phenomenon is also known to occur during cell-cycle arrest and is a characteristic of senescent cells (Marescal and Cheeseman 2020). Normally, proximal tubule cells *in vivo* predominantly do not undergo mitosis but re-enter the cell cycle after renal injury to repair tissue. Several animal studies have been published showing an increase in senescent cells upon drug induced renal injury with streptozotocin (STZ) and cisplatin as well as in mouse models of hypertension and diabetes (Liu et al. 2012; Westhoff et al. 2008; Zhou et al. 2004). Our data indicates that OTA is inducing a senescent-like phenotype in PTECs characterized by metabolic quiescence, which may hinder renal repair following injury; however, cell cycle and senescent-cell markers should be studied to confirm this. This study did not assess which components of mitochondrial oxidative phosphorylation (e.g. Complex I, II etc.) were inhibited by OTA, nor did we assess changes or disruption membrane potential. These are areas of future investigation to more finely delineate how OTA leads to mitochondrial toxicity.

We have previously shown that OTA downregulates numerous targets of the antioxidant response pathway regulated by NRF2, in accordance with many other studies (Boesch-Saadatmandi et al. 2009; Cavin et al. 2007; Imaoka et al. 2020). Here, we investigated whether the expression of these target genes can be recovered by directly activating NRF2 with two agonists, sulforaphane and tert-butylhydroquinone. tBHQ is a more potent agonist of NRF2 than sulforaphane; both act by hindering the ability of KEAP1 to sequester and ubiquitinate NRF2 in the cytosol (Houghton et al. 2016; Li and Kong 2009). OTA exposure resulted in over 7000 differentially expressed genes (FDR<0.05) compared with control in both donors. Surprisingly, co-treatment with SFN or tBHQ resulted in no differentially expressed genes compared with OTA treatment alone, except for 6 genes with tBHQ co-treatment in one donor (Table 2). Numerous NRF2 targets were downregulated in both donors, and neither SFN nor tBHQ significantly recovered expression in all cases (Figure 3A, C). However, PTECs did induce NRF2 target genes in response to either tBHQ or SFN (Figure 3B, PT6_LogCPM.csv and PT10_LogCPM.csv). This indicates that inhibiting KEAP1’s ability to bind and degrade NRF2 is not sufficient to recover oxidative response transcription and OTA likely does not hinder the liberation of NRF2 from KEAP1. More advanced techniques, such as fluorescence resonance energy transfer (FRET) imaging should be applied to confirm the binding and release mechanisms of NRF2 from KEAP1 in the tested exposure paradigms. Exploring other mechanisms, one study showed that OTA reduced the amount of phosphorylated NRF2 and NRF2 itself in the nucleus of mesenchymal stem cells, indicating that OTA may hinder nuclear translocation (Yoon et al. 2016). Pharmacological and dietary targeting of NRF2 has been examined as a prevention against OTA-mediated toxicity, with some success *in vitro* and in animal models. Ramyaa et al. showed that quercetin was able to alleviate OTA toxicity in HepG2 cells by reducing ROS and inducing NRF2-mediated expression (Ramyaa et al. 2014). Additionally, Liu et al. showed that Astaxanthin was also shown to upregulate NRF2 target expression and reduce OTA-induced cardiac injury in mice (Xu et al. 2019). Thus, the exact mechanism in which OTA suppresses the transcription of NRF2 target genes remains unknown.

Lastly, we investigated the role of inflammation due to OTA exposure, specifically by the cyclic-GMP-AMP synthase, stimulator of interferon genes (cGAS-STING) pathway activation. As we have shown evidence of OTA-induced mitochondrial damage in this study, we hypothesized that mtDNA released in the cytosol may activate cGAS-STING and lead to renal inflammation (Sun et al. 2023). Several downstream targets of type-1 interferons induced by STING were shown to be upregulated with OTA exposure in PT10 (Figure 4A). We chose to analyze protein levels of interleukin-6 (IL-6) and CXC motif chemokine-ligand 6 (CXCL6) due to their reported relevance to renal fibrosis and CKD (Ranganathan et al. 2013; Sun et al. 2019). We found that while 24-hour OTA exposure increased levels of IL-6 by 3-fold, co-administration of cGAS-STING inhibitor H-151 did not successfully ameliorate IL-6 increase in all three biological donors (Figure 4B). Interestingly, it did completely inhibit OTA-dependent IL-6 production in one donor (Supplementary Figure 2). OTA was shown to decrease CXCL6 levels slightly and further reduction was observed with co-administration of H-151, even though induction was not seen with positive control cGAMP treatment (Figure 4B). This is not surprising as there is little evidence of CXCL6 activation by the cGAS-STING pathway, despite its potential to be activated by type-1 interferons. Other groups have reported IL-6 induction with OTA in various animal cell lines of not only the kidney but microglia, gastric epithelium and liver (Chen et al. 2021; Raghubeer et al. 2017; Wang et al. 2019). One study found that 72-hour OTA exposure in human gastric endothelium cells (GES-1) induces pro-inflammatory cytokines by increased glycolysis and mTOR/HIF1α signaling (Wang et al. 2024). Evidence in this study is inconclusive in determining if STING activation is induced by OTA and is responsible for cytokine production, as it is likely that other pathways such as mTOR signaling are the major regulators. It should be noted that the present study did not test direct activation of cGAS-STING, but only a few downstream target mRNAs and protein levels of IL-6 with STING-specific modulators.

It is important to highlight the seemingly contradictory patterns in gene dysregulation observed in this report between the two donors used, PT6 and PT10. A limitation of this study is that only two biological donors were used in the mRNA sequencing experiment at a single timepoint of 72 hours, and differences observed in the transcriptome may be due to interindividual variability. With the nature of mRNA sequencing being a “snapshot” in time, and that differential gene expression does not always translate to altered cellular processes, the discrepancy between the two donors is not wholly unexpected. Notably, as listed in Supplementary Table 1, PT6 cells were isolated from a female donor and PT10 cells were isolated from a male donor, with the female donor having a history of heavy alcohol use and the male donor having a history of type 2 diabetes, which may explain the observed differences. For example, the upregulation of cytokines and chemokines reported in Figure 4A for PT10 may be confounded due to the history of diabetes in this donor. We also note that our primary PTECs did not exhibit measurable OAT1/3 (organic anion transporter 1/3) transcripts in RNA-seq, suggesting that OTA uptake under our culture conditions may occur via passive diffusion or alternative transporters at higher concentrations. Although OAT-mediated secretion may dominate OTA handling in vivo, these in vitro findings nonetheless reveal OTA’s intracellular actions on mitochondria and bioenergetics independent of OAT expression and the involvement of these transporters in OTA disposition in the renal straight tubule should be an area of future study. Nevertheless, biochemical and imaging data presented here show consistent, significant trends among three different donors, highlighting the importance of complementing transcriptional data with corresponding cellular and biochemical data.

In conclusion, the present study has shown that OTA induces mitochondrial dysfunction at a structural and functional level in biologically relevant primary human cells derived from the kidney proximal tubule at physiologically relevant exposure levels. The graphical abstract shows our hypothesized mechanism of OTA-dependent nephrotoxicity based on our findings. These findings are in support of mitochondrial dysfunction, renal hypoxia and oxidative stress as key toxicological mechanisms of OTA nephrotoxicity, as these pathways have been established as characteristic of chronic kidney disease progression. This study provides valuable insight into proximal-tubule specific pathways induced by OTA in humans in the context of renal disease. Chronic kidney disease of unknown etiology is thought to involve multifactorial environmental agents, and while this study does not provide a causative link between OTA and CKDu, it provides a better understanding of how OTA affects the kidney proximal tubule and is a potential contributing factor to CKDu.

## Supporting information

Supplementary Video S1-control

Supplementary Video S1-OTA

## CRediT authorship contribution statement

**Anish Mahadeo:** Investigation, formal analysis, writing – original draft, conceptualization. **Angela Zheng**: Investigation. **Edward Kelly:** Writing – review and editing, supervision, funding acquisition, resources. **Catherine Yeung:** Writing **–** review and editing, funding acquisition, supervision. **Theo Bammler:** Formal analysis, writing – review and editing. **James Macdonald:** Formal analysis, writing - review and editing. **Jonathan Himmelfarb**: Funding acquisition, writing – review and editing.

## Genomic Data Availability

Data is available on the Gene Expression Omnibus: GSE289962

## Acknowledgements

The authors would like to acknowledge Dr. Dale Hailey, University of Washington Institute of Stem Cells and Regenerative Medicine Lynn and Mike Garvey Imaging core director for assistance with confocal microscopy, supported by the Garvey family gift and the UW student technology fund. We would also like to thank Dr. Dennis Wang and Fu-Yen Chang, University of Washington Diabetes Research Center for assistance with Seahorse assay procedures and XFe96 Flux Analyzer usage, supported by the Diabetes Research Center Grant (NIH 5 P30 DK017047). We would also like to acknowledge Drs. Lans Taylor and Larry Vernetti for supplying the COX-8 GFP mitochondrial label vector. The graphical abstract was created in BioRender with license: Mahadeo, A. (2025) https://BioRender.com/ibeb54x.

---

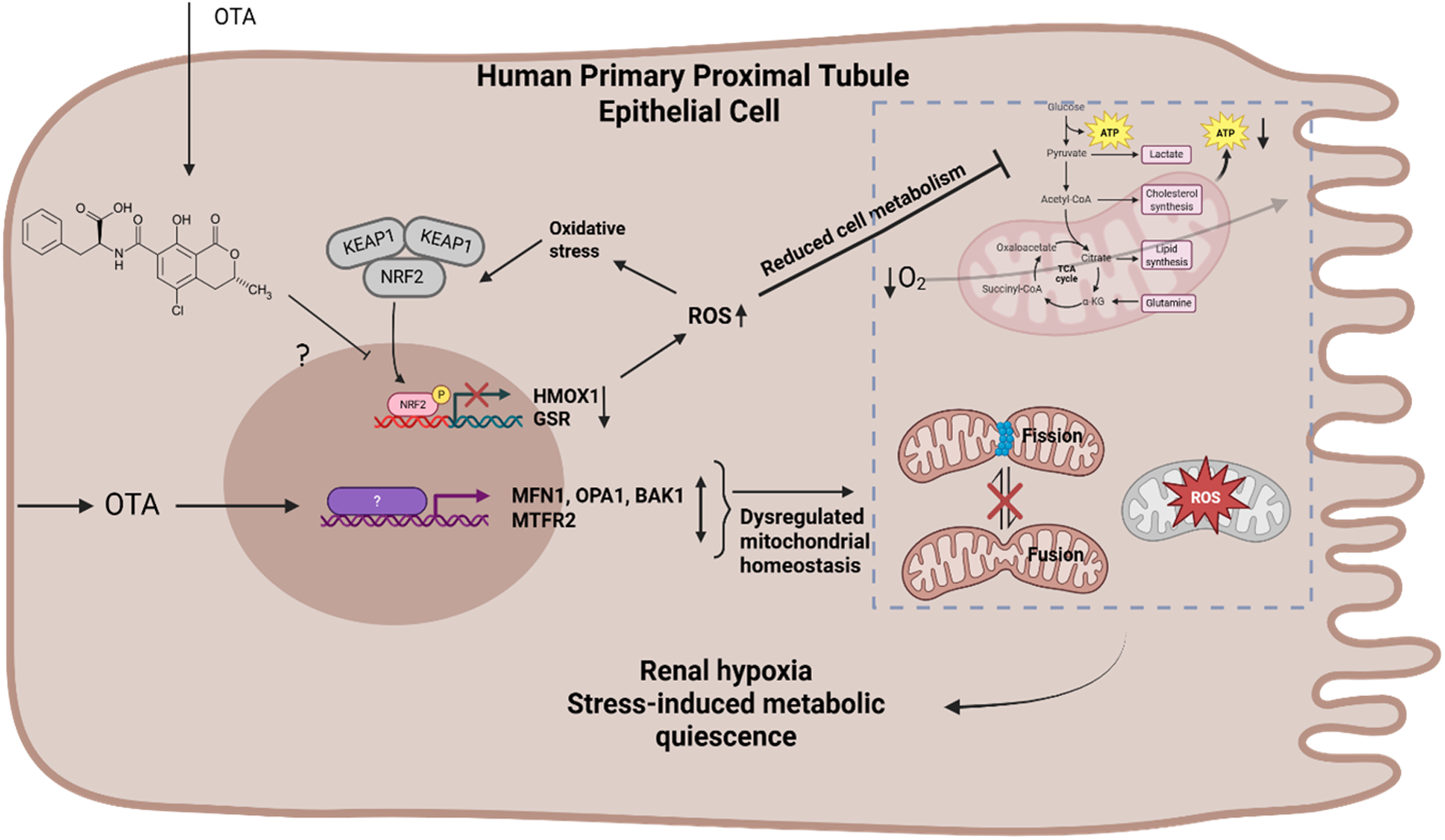

## References

Abid S et al. 2003. Ochratoxin a and human chronic nephropathy in tunisia: Is the situation endemic? Hum Exp Toxicol. 22(2):77–84. https://www.ncbi.nlm.nih.gov/pubmed/12693831. doi:10.1191/0960327103ht328oa.

Adebayo M, Singh S, Singh AP, Dasgupta S. 2021. Mitochondrial fusion and fission: The fine-tune balance for cellular homeostasis. FASEB J. 35(6):e21620. https://www.ncbi.nlm.nih.gov/pubmed/34048084. doi:10.1096/fj.202100067R.

Anzai N, Jutabha P, Endou H. 2010. Molecular mechanism of ochratoxin a transport in the kidney. Toxins (Basel). 2(6):1381–1398. https://www.ncbi.nlm.nih.gov/pubmed/22069643. doi:10.3390/toxins2061381.

Babayan N et al. 2020. Ochratoxin a induces global DNA hypomethylation and oxidative stress in neuronal cells in vitro. Mycotoxin Res. 36(1):73–81. https://www.ncbi.nlm.nih.gov/pubmed/31441013. doi:10.1007/s12550-019-00370-y.

Boesch-Saadatmandi C et al. 2009. Ochratoxin a impairs nrf2-dependent gene expression in porcine kidney tubulus cells. J Anim Physiol Anim Nutr (Berl). 93(5):547–554. https://www.ncbi.nlm.nih.gov/pubmed/18547363. doi:10.1111/j.1439-0396.2008.00838.x.

Cavin C et al. 2007. Reduction in antioxidant defenses may contribute to ochratoxin a toxicity and carcinogenicity. Toxicol Sci. 96(1):30–39. https://www.ncbi.nlm.nih.gov/pubmed/17110534. doi:10.1093/toxsci/kfl169.

Chalker B et al. 2025. Isolation of primary human proximal tubule epithelial cells and their use in creating a microphysiological model of the renal proximal tubule. J Vis Exp. 219). https://www.ncbi.nlm.nih.gov/pubmed/40418667. doi:10.3791/67997.

Chen Y et al. 2021. Astaxanthin alleviates ochratoxin a-induced cecum injury and inflammation in mice by regulating the diversity of cecal microbiota and tlr4/myd88/nf-kappab signaling pathway. Oxid Med Cell Longev. 2021(8894491. https://www.ncbi.nlm.nih.gov/pubmed/33505592. doi:10.1155/2021/8894491.

Chung KW et al. 2019. Mitochondrial damage and activation of the sting pathway lead to renal inflammation and fibrosis. Cell Metab. 30(4):784–799 e785. https://www.ncbi.nlm.nih.gov/pubmed/31474566. doi:10.1016/j.cmet.2019.08.003.

Cipolat S, Martins de Brito O, Dal Zilio B, Scorrano L. 2004. Opa1 requires mitofusin 1 to promote mitochondrial fusion. Proc Natl Acad Sci U S A. 101(45):15927–15932. https://www.ncbi.nlm.nih.gov/pubmed/15509649. doi:10.1073/pnas.0407043101.

Coronel MB, Sanchis V, Ramos AJ, Marin S. 2011. Ochratoxin a in adult population of lleida, spain: Presence in blood plasma and consumption in different regions and seasons. Food Chem Toxicol. 49(10):2697–2705. https://www.ncbi.nlm.nih.gov/pubmed/21802478. doi:10.1016/j.fct.2011.07.045.

Creppy EE, Roschenthaler R, Dirheimer G. 1984. Inhibition of protein synthesis in mice by ochratoxin a and its prevention by phenylalanine. Food Chem Toxicol. 22(11):883–886. https://www.ncbi.nlm.nih.gov/pubmed/6542055. doi:10.1016/0278-6915(84)90170-4.

de Groene EM, Jahn A, Horbach GJ, Fink-Gremmels J. 1996. Mutagenicity and genotoxicity of the mycotoxin ochratoxin a. Environ Toxicol Pharmacol. 1(1):21–26. https://www.ncbi.nlm.nih.gov/pubmed/21781659. doi:10.1016/1382-6689(95)00005-4.

Desalegn B et al. 2011. Mycotoxin detection in urine samples from patients with chronic kidney disease of uncertain etiology in sri lanka. Bull Environ Contam Toxicol. 87(1):6–10. https://www.ncbi.nlm.nih.gov/pubmed/21553028. doi:10.1007/s00128-011-0301-4.

Domijan AM et al. 1999. Ochratoxin a in blood of healthy population in zagreb. Arh Hig Rada Toksikol. 50(3):263–271. https://www.ncbi.nlm.nih.gov/pubmed/10649842.

Du Y, Gupta P, Qin S, Sieber M. 2023. The role of metabolism in cellular quiescence. J Cell Sci. 136(16). https://www.ncbi.nlm.nih.gov/pubmed/37589342. doi:10.1242/jcs.260787.

Edwards A, Kurtcuoglu V. 2022. Renal blood flow and oxygenation. Pflugers Arch. 474(8):759–770. https://www.ncbi.nlm.nih.gov/pubmed/35438336. doi:10.1007/s00424-022-02690-y.

Fazekas B, Tar A, Kovacs M. 2005. Ochratoxin a content of urine samples of healthy humans in hungary. Acta Vet Hung. 53(1):35–44. https://www.ncbi.nlm.nih.gov/pubmed/15782657. doi:10.1556/AVet.53.2005.1.4.

Fontecha-Barriuso M et al. 2022. Tubular mitochondrial dysfunction, oxidative stress, and progression of chronic kidney disease. Antioxidants (Basel). 11(7). https://www.ncbi.nlm.nih.gov/pubmed/35883847. doi:10.3390/antiox11071356.

Fuchs R, Peraica M. 2005. Ochratoxin a in human kidney diseases. Food Addit Contam. 22 Suppl 1(53–57. https://www.ncbi.nlm.nih.gov/pubmed/16332622. doi:10.1080/02652030500309368.

Gautier JC et al. 2001. Oxidative damage and stress response from ochratoxin a exposure in rats. Free Radic Biol Med. 30(10):1089–1098. https://www.ncbi.nlm.nih.gov/pubmed/11369498. doi:10.1016/s0891-5849(01)00507-x.

Gilkerson RW, Margineantu DH, Capaldi RA, Selker JM. 2000. Mitochondrial DNA depletion causes morphological changes in the mitochondrial reticulum of cultured human cells. FEBS Lett. 474(1):1–4. https://www.ncbi.nlm.nih.gov/pubmed/10828440. doi:10.1016/s0014-5793(00)01527-1.

Guo Q et al. 2024. Apelin regulates mitochondrial dynamics by inhibiting mst1-jnk-drp1 signaling pathway to reduce neuronal apoptosis after spinal cord injury. Neurochem Int. 180(105885. https://www.ncbi.nlm.nih.gov/pubmed/39433147. doi:10.1016/j.neuint.2024.105885.

He X, Ma Q. 2009. Nrf2 cysteine residues are critical for oxidant/electrophile-sensing, kelch-like ech-associated protein-1-dependent ubiquitination-proteasomal degradation, and transcription activation. Mol Pharmacol. 76(6):1265–1278. https://www.ncbi.nlm.nih.gov/pubmed/19786557. doi:10.1124/mol.109.058453.

Houghton CA, Fassett RG, Coombes JS. 2016. Sulforaphane and other nutrigenomic nrf2 activators: Can the clinician’s expectation be matched by the reality? Oxid Med Cell Longev. 2016(7857186. https://www.ncbi.nlm.nih.gov/pubmed/26881038. doi:10.1155/2016/7857186.

Imaoka T et al. 2020. Microphysiological system modeling of ochratoxin a-associated nephrotoxicity. Toxicology. 444(152582. https://www.ncbi.nlm.nih.gov/pubmed/32905824. doi:10.1016/j.tox.2020.152582.

Jayasekara KB et al. 2019. Relevance of heat stress and dehydration to chronic kidney disease (ckdu) in sri lanka. Prev Med Rep. 15(100928. https://www.ncbi.nlm.nih.gov/pubmed/31304082. doi:10.1016/j.pmedr.2019.100928.

Jutabha P et al. 2011. A novel human organic anion transporter npt4 mediates the transport of ochratoxin a. J Pharmacol Sci. 116(4):392–396. https://www.ncbi.nlm.nih.gov/pubmed/21778665. doi:10.1254/jphs.10227sc.

Karbowski M, Youle RJ. 2003. Dynamics of mitochondrial morphology in healthy cells and during apoptosis. Cell Death Differ. 10(8):870–880. https://www.ncbi.nlm.nih.gov/pubmed/12867994. doi:10.1038/sj.cdd.4401260.

Kim D, Paggi JM, Park C, Bennett C, Salzberg SL. 2019. Graph-based genome alignment and genotyping with hisat2 and hisat-genotype. Nat Biotechnol. 37(8):907–915. https://www.ncbi.nlm.nih.gov/pubmed/31375807. doi:10.1038/s41587-019-0201-4.

Kosicki R, Buharowska-Donten J, Twaruzek M. 2021. Ochratoxin a levels in serum of polish dialysis patients with chronic renal failure. Toxicon. 200(183-188. https://www.ncbi.nlm.nih.gov/pubmed/34375657. doi:10.1016/j.toxicon.2021.08.002.

Law CW, Chen Y, Shi W, Smyth GK. 2014. Voom: Precision weights unlock linear model analysis tools for rna-seq read counts. Genome Biol. 15(2):R29. https://www.ncbi.nlm.nih.gov/pubmed/24485249. doi:10.1186/gb-2014-15-2-r29.

Legros F, Lombes A, Frachon P, Rojo M. 2002. Mitochondrial fusion in human cells is efficient, requires the inner membrane potential, and is mediated by mitofusins. Mol Biol Cell. 13(12):4343–4354. https://www.ncbi.nlm.nih.gov/pubmed/12475957. doi:10.1091/mbc.e02-06-0330.

Li W, Kong AN. 2009. Molecular mechanisms of nrf2-mediated antioxidant response. Mol Carcinog. 48(2):91–104. https://www.ncbi.nlm.nih.gov/pubmed/18618599. doi:10.1002/mc.20465.

Liao Y, Smyth GK, Shi W. 2019. The r package rsubread is easier, faster, cheaper and better for alignment and quantification of rna sequencing reads. Nucleic Acids Res. 47(8):e47. https://www.ncbi.nlm.nih.gov/pubmed/30783653. doi:10.1093/nar/gkz114.

Limonciel A, Jennings P. 2014. A review of the evidence that ochratoxin a is an nrf2 inhibitor: Implications for nephrotoxicity and renal carcinogenicity. Toxins (Basel). 6(1):371–379. https://www.ncbi.nlm.nih.gov/pubmed/24448208. doi:10.3390/toxins6010371.

Liu J et al. 2012. Accelerated senescence of renal tubular epithelial cells is associated with disease progression of patients with immunoglobulin a (iga) nephropathy. Transl Res. 159(6):454–463. https://www.ncbi.nlm.nih.gov/pubmed/22633096. doi:10.1016/j.trsl.2011.11.008.

Loboda A et al. 2017. Nrf2 deficiency exacerbates ochratoxin a-induced toxicity in vitro and in vivo. Toxicology. 389(42–52. https://www.ncbi.nlm.nih.gov/pubmed/28710020. doi:10.1016/j.tox.2017.07.004.

Loor G et al. 2010. Menadione triggers cell death through ros-dependent mechanisms involving parp activation without requiring apoptosis. Free Radic Biol Med. 49(12):1925–1936. https://www.ncbi.nlm.nih.gov/pubmed/20937380. doi:10.1016/j.freeradbiomed.2010.09.021.

Maaroufi K et al. 1995. Ochratoxin a in human blood in relation to nephropathy in tunisia. Hum Exp Toxicol. 14(7):609–614. https://www.ncbi.nlm.nih.gov/pubmed/7576823. doi:10.1177/096032719501400710.

Malir F, Ostry V, Pfohl-Leszkowicz A, Malir J, Toman J. 2016. Ochratoxin a: 50 years of research. Toxins (Basel). 8(7). https://www.ncbi.nlm.nih.gov/pubmed/27384585. doi:10.3390/toxins8070191.

Mantle PG, Faucet-Marquis V, Manderville RA, Squillaci B, Pfohl-Leszkowicz A. 2010. Structures of covalent adducts between DNA and ochratoxin a: A new factor in debate about genotoxicity and human risk assessment. Chem Res Toxicol. 23(1):89–98. https://www.ncbi.nlm.nih.gov/pubmed/19928877. doi:10.1021/tx900295a.

Marescal O, Cheeseman IM. 2020. Cellular mechanisms and regulation of quiescence. Dev Cell. 55(3):259–271. https://www.ncbi.nlm.nih.gov/pubmed/33171109. doi:10.1016/j.devcel.2020.09.029.

McCarthy DJ, Smyth GK. 2009. Testing significance relative to a fold-change threshold is a treat. Bioinformatics. 25(6):765–771. https://www.ncbi.nlm.nih.gov/pubmed/19176553. doi:10.1093/bioinformatics/btp053.

O’Mealey GB, Berry WL, Plafker SM. 2017. Sulforaphane is a nrf2-independent inhibitor of mitochondrial fission. Redox Biol. 11(103-110. <GO to ISI>://WOS:000398212000011. doi:10.1016/j.redox.2016.11.007.

Paidi G et al. 2021. Chronic kidney disease of unknown origin: A mysterious epidemic. Cureus. 13(8):e17132. https://www.ncbi.nlm.nih.gov/pubmed/34548965. doi:10.7759/cureus.17132.

Palli D et al. 1999. Serum levels of ochratoxin a in healthy adults in tuscany: Correlation with individual characteristics and between repeat measurements. Cancer Epidemiol Biomarkers Prev. 8(3):265–269. https://www.ncbi.nlm.nih.gov/pubmed/10090305.

Park S, Lim W, You S, Song G. 2019. Ochratoxin a exerts neurotoxicity in human astrocytes through mitochondria-dependent apoptosis and intracellular calcium overload. Toxicol Lett. 313(42–49. https://www.ncbi.nlm.nih.gov/pubmed/31154016. doi:10.1016/j.toxlet.2019.05.021.

Raghubeer S, Nagiah S, Chuturgoon AA. 2017. Acute ochratoxin a exposure induces inflammation and apoptosis in human embryonic kidney (hek293) cells. Toxicon. 137(48–53. https://www.ncbi.nlm.nih.gov/pubmed/28712913. doi:10.1016/j.toxicon.2017.07.013.

Rajapaksha H, Pandithavidana DR, Dahanayake JN. 2021. Demystifying chronic kidney disease of unknown etiology (ckdu): Computational interaction analysis of pesticides and metabolites with vital renal enzymes. Biomolecules. 11(2). https://www.ncbi.nlm.nih.gov/pubmed/33578980. doi:10.3390/biom11020261.

Ramyaa P, Krishnaswamy R, Padma VV. 2014. Quercetin modulates ota-induced oxidative stress and redox signalling in hepg2 cells - up regulation of nrf2 expression and down regulation of nf-kappab and cox-2. Biochim Biophys Acta. 1840(1):681–692. https://www.ncbi.nlm.nih.gov/pubmed/24161694. doi:10.1016/j.bbagen.2013.10.024.

Ramyaa P, Padma VV. 2013. Ochratoxin-induced toxicity, oxidative stress and apoptosis ameliorated by quercetin--modulation by nrf2. Food Chem Toxicol. 62(205–216. https://www.ncbi.nlm.nih.gov/pubmed/23994659. doi:10.1016/j.fct.2013.08.048.

Ranganathan P, Jayakumar C, Ramesh G. 2013. Proximal tubule-specific overexpression of netrin-1 suppresses acute kidney injury-induced interstitial fibrosis and glomerulosclerosis through suppression of il-6/stat3 signaling. Am J Physiol Renal Physiol. 304(8):F1054–1065. https://www.ncbi.nlm.nih.gov/pubmed/23408169. doi:10.1152/ajprenal.00650.2012.

Schaaf GJ et al. 2002. The role of oxidative stress in the ochratoxin a-mediated toxicity in proximal tubular cells. Biochim Biophys Acta. 1588(2):149–158. https://www.ncbi.nlm.nih.gov/pubmed/12385779. doi:10.1016/s0925-4439(02)00159-x.

Schwerdt G, Bauer K, Gekle M, Silbernagl S. 1996. Accumulation of ochratoxin a in rat kidney in vivo and in cultivated renal epithelial cells in vitro. Toxicology. 114(3):177–185. https://www.ncbi.nlm.nih.gov/pubmed/8980707. doi:10.1016/s0300-483x(96)03484-1.

Shin HS, Lee HJ, Pyo MC, Ryu D, Lee KW. 2019. Ochratoxin a-induced hepatotoxicity through phase i and phase ii reactions regulated by ahr in liver cells. Toxins. 11(7). <GO to ISI>://WOS:000482110000051. doi:ARTN 377 10.3390/toxins11070377.

Skulachev VP. 2001. Mitochondrial filaments and clusters as intracellular power-transmitting cables. Trends Biochem Sci. 26(1):23–29. https://www.ncbi.nlm.nih.gov/pubmed/11165513. doi:10.1016/s0968-0004(00)01735-7.

Studer-Rohr I, Schlatter J, Dietrich DR. 2000. Kinetic parameters and intraindividual fluctuations of ochratoxin a plasma levels in humans. Arch Toxicol. 74(9):499–510. https://www.ncbi.nlm.nih.gov/pubmed/11131029. doi:10.1007/s002040000157.

Sun C et al. 2023. The activation of cgas-sting in acute kidney injury. J Inflamm Res. 16(4461–4470. https://www.ncbi.nlm.nih.gov/pubmed/37842189. doi:10.2147/JIR.S423232.

Sun MY et al. 2019. Cxcl6 promotes renal interstitial fibrosis in diabetic nephropathy by activating jak/stat3 signaling pathway. Front Pharmacol. 10(224. https://www.ncbi.nlm.nih.gov/pubmed/30967776. doi:10.3389/fphar.2019.00224.

Tagaya M et al. 2022. Inhibition of mitochondrial fission protects podocytes from albumin-induced cell damage in diabetic kidney disease. Biochim Biophys Acta Mol Basis Dis. 1868(5):166368. https://www.ncbi.nlm.nih.gov/pubmed/35202791. doi:10.1016/j.bbadis.2022.166368.

van der Reest J, Lilla S, Zheng L, Zanivan S, Gottlieb E. 2018. Proteome-wide analysis of cysteine oxidation reveals metabolic sensitivity to redox stress. Nat Commun. 9(1):1581. https://www.ncbi.nlm.nih.gov/pubmed/29679077. doi:10.1038/s41467-018-04003-3.

Van Ness KP et al. 2017. Microphysiological systems to assess nonclinical toxicity. Curr Protoc Toxicol. 73(14 18 11–14 18 28. https://www.ncbi.nlm.nih.gov/pubmed/28777442. doi:10.1002/cptx.27.

Wang W et al. 2019. Ochratoxin a induces liver inflammation: Involvement of intestinal microbiota. Microbiome. 7(1):151. https://www.ncbi.nlm.nih.gov/pubmed/31779704. doi:10.1186/s40168-019-0761-z.

Wang Y et al. 2024. Ochratoxin a-enhanced glycolysis induces inflammatory responses in human gastric epithelium cells through mtor/hif-1alpha signaling pathway. Ecotoxicol Environ Saf. 270(115868. https://www.ncbi.nlm.nih.gov/pubmed/38142590. doi:10.1016/j.ecoenv.2023.115868.

Weber EJ et al. 2016. Development of a microphysiological model of human kidney proximal tubule function. Kidney Int. 90(3):627–637. https://www.ncbi.nlm.nih.gov/pubmed/27521113. doi:10.1016/j.kint.2016.06.011.

Westhoff JH et al. 2008. Hypertension induces somatic cellular senescence in rats and humans by induction of cell cycle inhibitor p16ink4a. Hypertension. 52(1):123–129. https://www.ncbi.nlm.nih.gov/pubmed/18504326. doi:10.1161/HYPERTENSIONAHA.107.099432.

Xu W et al. 2019. Astaxanthin protects ota-induced lung injury in mice through the nrf2/nf-kappab pathway. Toxins (Basel). 11(9). https://www.ncbi.nlm.nih.gov/pubmed/31533259. doi:10.3390/toxins11090540.

Yoon DS, Choi Y, Lee JW. 2016. Cellular localization of nrf2 determines the self-renewal and osteogenic differentiation potential of human mscs via the p53-sirt1 axis. Cell Death Dis. 7(2):e2093. https://www.ncbi.nlm.nih.gov/pubmed/26866273. doi:10.1038/cddis.2016.3.

Yoon YS et al. 2006. Formation of elongated giant mitochondria in dfo-induced cellular senescence: Involvement of enhanced fusion process through modulation of fis1. J Cell Physiol. 209(2):468–80. https://www.ncbi.nlm.nih.gov/pubmed/16883569. doi:10.1002/jcp.20753.

Zhang X, Agborbesong E, Li X. 2021. The role of mitochondria in acute kidney injury and chronic kidney disease and its therapeutic potential. Int J Mol Sci. 22(20). https://www.ncbi.nlm.nih.gov/pubmed/34681922. doi:10.3390/ijms222011253.

Zhao C et al. 2017. Pink1/parkin-mediated mitophagy play a protective role in cisplatin induced renal tubular epithelial cells injury. Exp Cell Res. 350(2):390–397. https://www.ncbi.nlm.nih.gov/pubmed/28024839. doi:10.1016/j.yexcr.2016.12.015.

Zheng F, Chen P, Li H, Aschner M. 2020. Drp-1-dependent mitochondrial fragmentation contributes to cobalt chloride-induced toxicity in caenorhabditis elegans. Toxicol Sci. 177(1):158–167. https://www.ncbi.nlm.nih.gov/pubmed/32617571. doi:10.1093/toxsci/kfaa105.

Zhou H et al. 2004. The induction of cell cycle regulatory and DNA repair proteins in cisplatin-induced acute renal failure. Toxicol Appl Pharmacol. 200(2):111–120. https://www.ncbi.nlm.nih.gov/pubmed/15476864. doi:10.1016/j.taap.2004.04.003.

