## Supplementary Materials for "Evaluation of toxicological mechanisms of ochratoxin-A in human primary proximal tubule epithelial cells"

| **Table S1. Donor information on kidney tissues from which PTECs were isolated.** | | | | | |
| --- | --- | --- | --- | --- | --- |
| **Donor ID** | **Age** | **Sex** | **Ethnicity** | **Pre-existing conditions** | **Final Pathology** |
| PT10 | 49 | Male | White | Diabetes, hypertension, chronic anemia, prior drug use | Cause of death: cardiovascular anoxia |
| PT11 | 65 | Male | White | Diabetes, hypertension, coronary artery disease, history of cigarette use | Cause of death: stroke, intracranial hemorrhage |
| PT6 | 45 | Female | White | Hypertension, history of heavy alcohol use | Cause of death: cerebrovascular stroke |
| PT21 | 52 | Female | White |  | Cause of death: cerebrovascular stroke |
| PT23 | 73 | Male | White | Diabetes, hypertension | Cause of death: Natural causes, anoxia |
| PT16 | 59 | Male | White | - | Cause of death: head trauma/blunt injury |
| PT3 | 42 | Female | White | History of opioid use | Cause of death: opioid overdose |

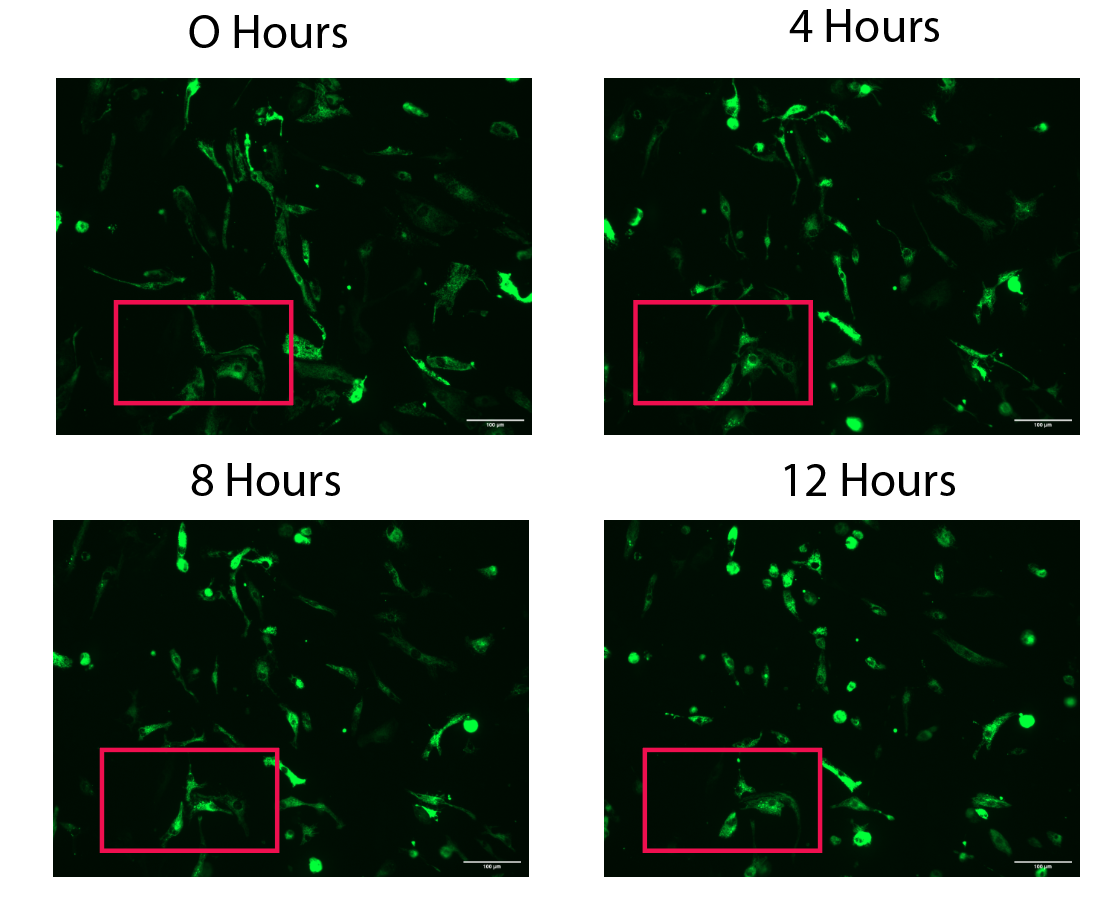

**Figure S1.** 12-hour time-lapse imaging of mitochondrial membrane-GFP labeled PTECs (PT23) treated with 10 μM OTA. The indicated region of interest (ROI) depicts mitochondria hyperfusion over time. Scale bar: 100 μm. Associated files: OTA-timelapse.avi, OTA_ROI.avi (indicated region), control_timelapse.avi

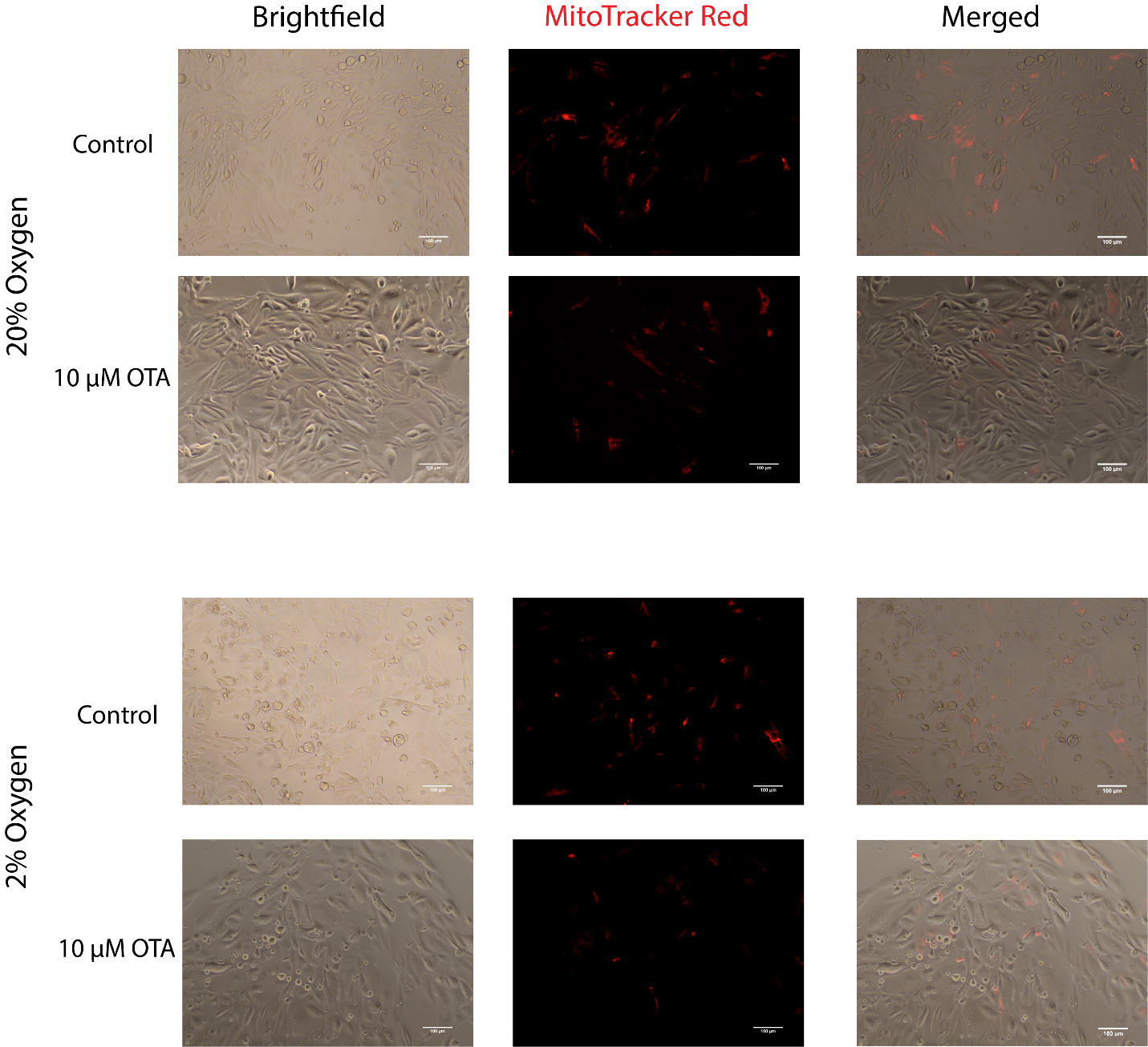

**Figure S2.** Fluorescence microscopy of mitochondria in PTECs (PT11) under normal oxygen conditions or renal cortex oxygen conditions (2%) after 24 hours. Mitochondrial signal reduction was observed with OTA exposure under 2% oxygen and to a lesser extent under 20% oxygen. Scale bar: 100 μm.

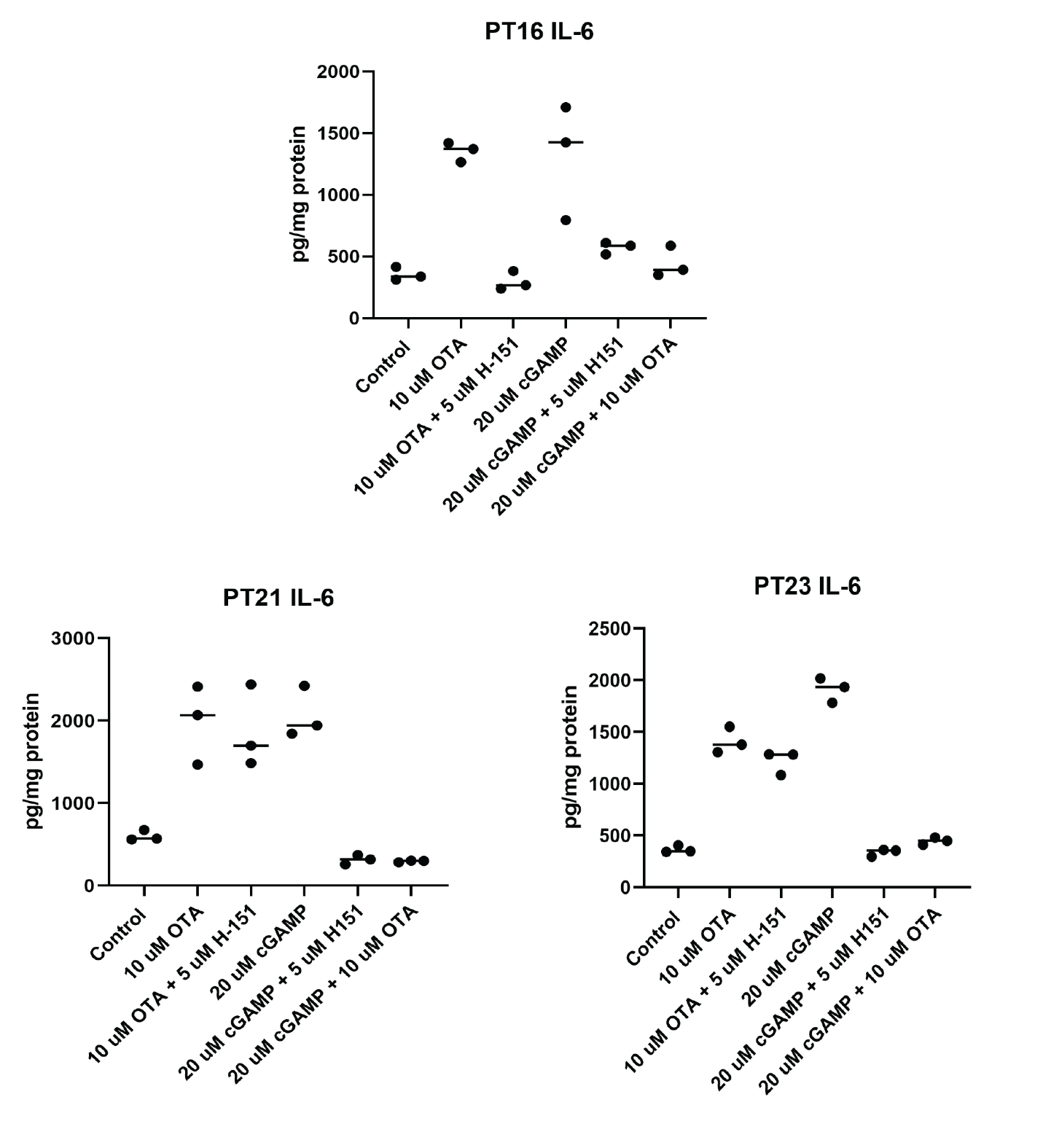

**Figure S3.** IL-6 ELISA data from Figure 4B separated by individual cell donor. Notably in PT16, OTA-induction of IL-6 levels was inhibited by co-administration of the STING inhibitor H-151.

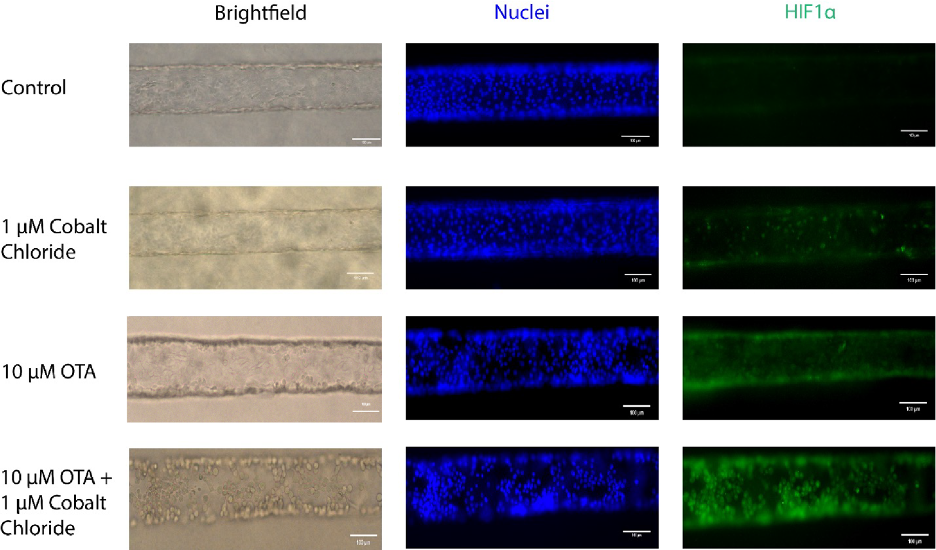

**Figure S4.** Brightfield and fluorescence microscopy reveals substantial nuclear stabilization of HIF1α in a proximal tubule microphysiological system (Donor: PT10) under co-administration of OTA and cobalt chloride. Images were taken after 48 hours of exposure. Scale bar: 100 μm.
